# Uncovered genetic diversity in *Hemidactylus mabouia* (Reptilia: Gekkonidae) from Madeira Island reveals uncertain sources of introduction

**DOI:** 10.1101/2021.03.02.433159

**Authors:** Catarina Rato, Beatriz Martins, Ricardo Rocha, Iolanda Silva-Rocha

**Author notes:** These authors have contributed equally to this work (Ricardo Rocha, Iolanda Silva-Rocha).

## Abstract

*Hemidactylus mabouia* is one of the most widely distributed species within its genus. It was first reported to Madeira in 2002 and the first individuals were considered to have originated from Cape Verde. Almost 20 years later, we found that *H. mabouia* has substantially expanded its distribution and can now be found >10 km away from Funchal, where it was first reported. Based on a 12S phylogenetic analysis and using 29 individuals from Funchal and Câmara de Lobos we found that Madeira actually harbours two distinct lineages of *H. mabouia*: one exclusively South American and another widespread in America and Africa. However, the lack of genetic diversity typical of this species outside its native range and the obtained phylogenetic pattern prevent us to infer possible introduction routes or sources. Our study emphasizes that authorities should remain vigilant regarding the arrival of other non-natives and act to prevent their establishment as soon as they are detected.

---

The rate of species’ translocation to areas outside their native geographical range is increasing with human globalization, considerably impacting native biotas (Simberloff et al, 2013). Reptiles are particularly prone to biological invasions, either by being one of the most often introduced animal groups, or by being notably sensitive to the impacts of invaders (Kraus, 2009). The introduction of species to island ecosystems is particularly problematic, due to the limited capacity of insular native species to compete with newcomers, elude predators and handle diseases, inherent from their evolutionary insular history (Novosolov et al, 2013; Whittaker and Fernández-Palacios, 2007). *Hemidactylus* Gray, 1845 is one of the most species rich genera within the Gekkonidae family, currently comprising about 164 recognized species (Uetz et al, 2020). These geckos have their native distribution throughout most of tropical Asia and Africa, the more arid regions of northeast Africa and southwest Asia, the Mediterranean region and South America, which they reached by natural transmarine colonization (Kluge, 1969). They have recently colonized, both naturally and anthropogenically, other areas of the Americas and the West Indies, Australia and several islands of the Indian, Pacific and Atlantic Oceans (see references in Carranza and Arnold, 2006). Their success as human-assisted colonizers is partly associated with the synanthropic habits of several *Hemidactylus* species, allied with their relatively small size and cryptic nature (Hoskin, 2011). The tropical house gecko *Hemidactylus mabouia* is one of the most successful invaders within the genus, and it can now be found in both the Old and New Worlds, as well as in many Atlantic islands, such as Cape Verde (Jesus et al, 2001; although classified later as *Hemidactylus mercatorius* in Vasconcelos et al, 2013 following Rocha et al, 2010), the Gulf of Guinea Islands (Jesus et al, 2005) and Madeira (Jesus et al, 2002). The successive introductions and absence of genetic differentiation within *H. mabouia* over its extensive range, precludes the assessment of the ultimate geographic origin of this species (Carranza and Arnold, 2006). Nonetheless, the great genetic diversity observed on Mayotte island in the Comoros, and the relationship to northeast Madagascar populations assigned to *H. mercatorius*, suggest that the widespread haplotypes originated in East Africa, and from there spread westwards, across tropical Africa reaching several Atlantic islands and large parts of tropical America (Carranza and Arnold, 2006). Additionally, based on the deep divergence observed between African and Malagasy specimens, Vences et al (2004) began to refer to all *H. mabouia* populations in Madagascar as *Hemidactylus mercatorius* (Gray, 1842). Moreover, results from Rocha et al (2005) clearly show that some specimens of *H. mabouia* from the Gulf of Guinea cluster with one *H. mercatorius* clade from northern Madagascar and the Comoros. Because Boumans et al (2007) argued in favour of the native status of *H. mercatorius* across Madagascar, Rocha et al (2010) proposed that insular populations of *H. mabouia* should be recognized as *H. mercatorius*, and the status of East African populations should remain unchanged as *H. mabouia*, pending further studies. Clearly, the complex phylogeographical patterns and taxonomy of the putative “species-complex” *H. mabouia/mercatorius* is far from being understood.

In this study, we focus on the genetic affinities of *H. mabouia* specimens from Madeira Island, in an attempt to infer the geographical origin of the introduced populations. The volcanic archipelago of Madeira is located ca. 520 km from the African coast and is comprised by the islands of Madeira, Porto Santo and Desertas (fig. 1). Almost 20 years ago, Jesus et al (2002) reported the first three specimens in Madeira, captured in the island’s capital Funchal. These animals were assigned as *H. mabouia* based on their genetic similarity with other sequences available at the time on GenBank (12S and cytb). Through the analysis by eye of the alignment, the authors concluded that the Madeiran specimens were most similar to previously sampled individuals from the Cape Verdean island of São Vicente. Here, apart from collecting specimens from an additional locality in Madeira, the total number of analysed individuals is substantially larger, allowing the detection of possible within island intraspecific genetic variation. Additionally, through a phylogenetic analysis, we use available *H. mabouia* sequences from the species’ known distribution to infer possible routes and sources of introduction of Madeira’s populations of tropical house geckos.

**Figure 1.**
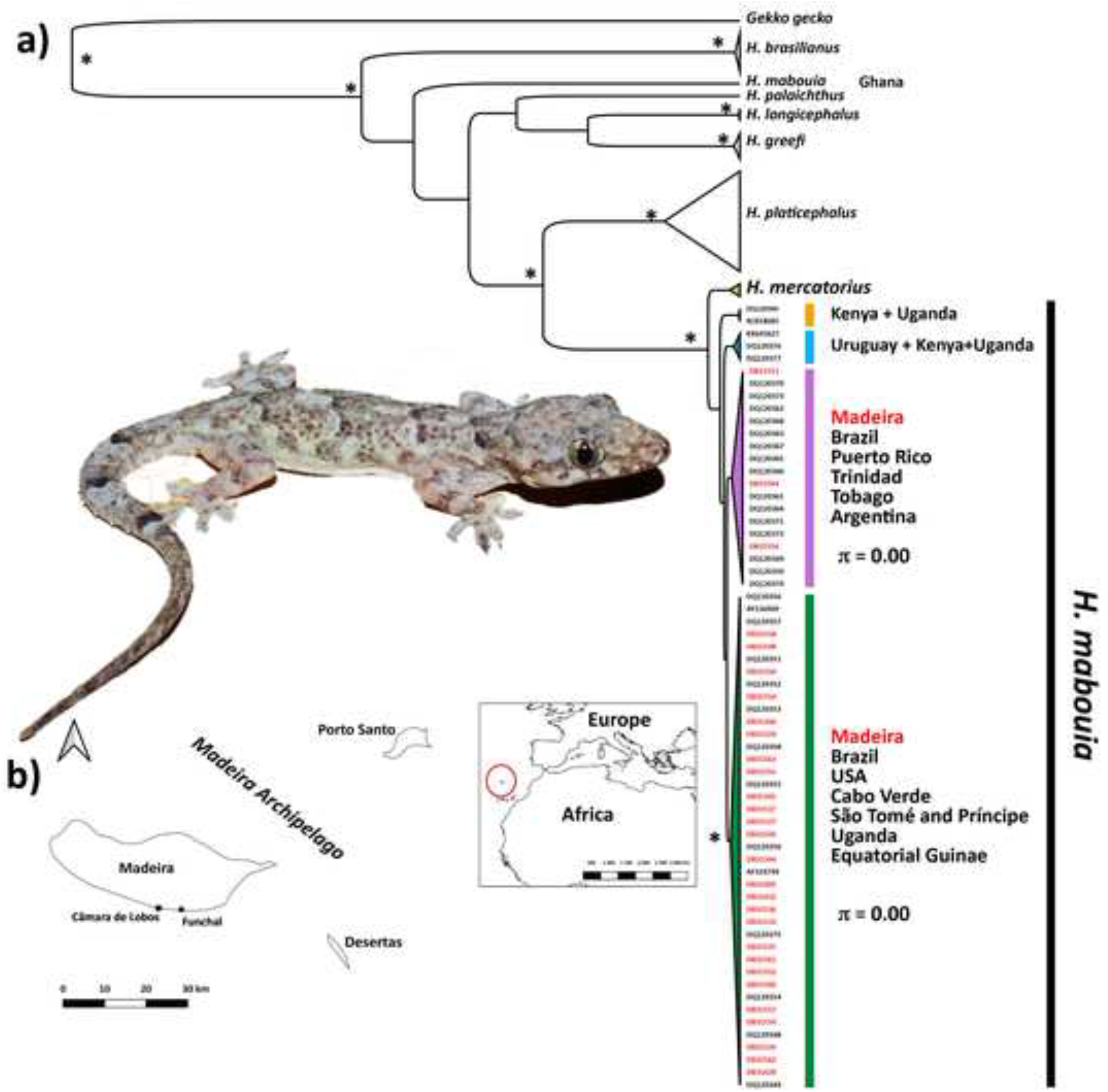
a) Bayesian phylogenetic tree derived from the 12S rRNA fragment and using *Gekko gecko* as outgroup. The stars next to the nodes represent values of posterior probabilities higher than 95%. The symbol π stands for genetic diversity; b) map depicting the geographic position of the Madeira Archipelago on a global scale, and in more detail the location of the two sampled populations (Funchal and Câmara de Lobos) in Madeira Island.

A total of 29 individuals assigned in the field to *H. mabouia* were collected from two distinct localities in Madeira, namely Funchal (N=16) and Câmara de Lobos (N=13). In Câmara de Lobos, some individuals were collected in the city centre, while others captured in a nearby industrial area.

Tissue from tail tip muscle was collected from each individual and preserved in 96% ethanol. Genomic DNA was extracted using an Easy-Spin Kit (Citomed). A fragment of the 12S rRNA gene was amplified by Polymerase Chain Reaction (PCR) using the primers L1091 and H1478 published by Kocher et al (1989). Amplification of the 12S mtDNA fragment was carried out in a 25 μl volume, containing 0.16 μM of each primer, 1U of Taq DNA polymerase (MyTaq HS Mix, Bioline), and approximately 100 ng of template DNA. PCR consisted of an initial denaturation at 95°C for 10 min, followed by 40 cycles that included a denaturation step at 95°C for 30 sec, annealing at 54°C for 30 sec and extension at 72°C for 30 sec. A final extension was conducted for 10 min. All successful amplifications were sent to CTM (CIBIO-InBIO) for purification and Sanger sequencing. This same protocol was also employed on two *H. mercatorius* specimens from Madagascar, with codes ACP1143 (Sahamalaza) and ACP2350 (Isalo).

A total of 35 12S sequences of *H. mabouia* were retrieved from GenBank and added to the dataset. Geographic location of each population from Madeira is depicted in fig. 1, and detailed information about the locality and GenBank accession codes is presented in Table 1. GenBank accession codes for the used *H. mercatorius* are MW665155 and MW665156. The obtained sequences were imported into the software Geneious Prime® (v.2020.2.4 Biomatters Ltd.) where the alignment was performed using MAFFT v.7.017 (Katoh and Standley, 2013) under the default parameters. Phylogenetic analysis based on the 12S mitochondrial fragment was performed under a Bayesian Inference (BI) method, using *Gekko gecko* as the outgroup, and several other representatives of the genus *Hemidactylus*, such as *H. mercatorius*, *H. longicephalus*, *H. greefi*, *H. brasilianus*, *H. palaichthus* and *H. platycephalus*, known to be closely related to *H. mabouia* (Carranza and Arnold, 2006; Gamble et al, 2011). In order to determine the best fitting nucleotide model, we used the software PartitionFinder v.2 (Lanfear et al, 2017), considering *branchlengths* = linked and *modelselection* = BIC. The software BEAST v.1.8.4 (Drummond and Rambaut, 2007) was used for the BI topology. Analyses were run twice for 25×10^6^ generations with a sampling frequency of 100. Models and prior specifications applied were as follows (otherwise by default): Strict Clock and Coalescent with Constant Population Size. Convergence for all model parameters was assessed by examining trace plots and histograms in Tracer v.1.7 (Rambaut et al, 2018) after obtaining an effective sample size (ESS) > 200. The initial 10% of samples were discarded as burn-in. Runs were combined using LogCombiner, and maximum credibility trees with divergence time means and 95% highest probability densities (HPDs) were produced using Tree Annotator. Trees were visualized using FigTree v.1.4.0 (Rambaut, 2009).

**Table 1.** Information regarding the specimens of *Hemidactylus mabouia* from Madeira used in this study.

| Specimen Code | Locality | Geographic coordinates | GenBank Accession Code |
| --- | --- | --- | --- |
| DB31344 | Funchal | 32.644349, -16.915567 | MW665176 |
| DB31345 | Funchal | 32.644349, -16.915567 | MW665178 |
| DB31346 | Funchal | 32.644349, -16.915567 | MW665180 |
| DB31405 | Funchal | 32.644349, -16.915567 | MW665161 |
| DB31428 | Funchal | 32.644349, -16.915567 | MW665182 |
| DB31432 | Funchal | 32.644349, -16.915567 | MW665166 |
| DB31524 | Funchal | 32.644349, -16.915567 | MW665169 |
| DB31525 | Funchal | 32.644349, -16.915567 | MW665173 |
| DB31526 | Funchal | 32.644349, -16.915567 | MW665177 |
| DB31527 | Funchal | 32.644349, -16.915567 | MW665179 |
| DB31533 | Funchal | 32.644349, -16.915567 | MW665181 |
| DB31536 | Funchal | 32.644349, -16.915567 | MW665162 |
| DB31537 | Funchal | 32.644349, -16.915567 | MW665165 |
| DB31548 | Funchal | 32.644349, -16.915567 | MW665184 |
| DB31549 | Funchal | 32.644349, -16.915567 | MW665170 |
| DB31550 | Funchal | 32.644349, -16.915567 | MW665174 |
| DB31553 | Câmara de Lobos (village) | 32.647043, -16.973176 | MW665157 |
| DB31559 | Câmara de Lobos (village) | 32.647043, -16.973176 | MW665163 |
| DB31564 | Câmara de Lobos (village) | 32.647043, -16.973176 | MW665158 |
| DB31552 | Câmara de Lobos (industrial) | 32.647220, -16.969386 | MW665167 |
| DB31554 | Câmara de Lobos (industrial) | 32.647220, -16.969386 | MW665171 |
| DB31555 | Câmara de Lobos (industrial) | 32.647220, -16.969386 | MW665175 |
| DB31556 | Câmara de Lobos (industrial) | 32.647220, -16.969386 | MW665159 |
| DB31557 | Câmara de Lobos (industrial) | 32.647220, -16.969386 | MW665185 |
| DB31558 | Câmara de Lobos (industrial) | 32.647220, -16.969386 | MW665160 |
| DB31560 | Câmara de Lobos (industrial) | 32.647220, -16.969386 | MW665164 |
| DB31561 | Câmara de Lobos (industrial) | 32.647220, -16.969386 | MW665183 |
| DB31562 | Câmara de Lobos (industrial) | 32.647220, -16.969386 | MW665168 |
| DB31563 | Câmara de Lobos (industrial) | 32.647220, -16.969386 | MW665172 |

According to the obtained 12S phylogenetic topology, the genetic diversity of *H. mabouia* is represented by four clades, with *H. mercatorius* as its sister taxon (fig. 1). Nevertheless, most of these relationships are not highly supported. While this pattern within *H. mabouia* is not a novelty (see Carranza and Arnold, 2006), the genetic diversity found in Madeira is. Surprisingly, apart from the specimens already identified by Jesus et al (2002) as related to individuals of *H. mabouia* from Cape Verde, this study reveals the existence of three individuals belonging to a new lineage in Madeira, related exclusively to South American specimens. Only individuals captured in Câmara de Lobos, both in the city and industrial area, where assigned to this new lineage. Though Câmara de Lobos is mostly a traditional fishing town with no international harbour, its industrial park is one of the island’s most active commercial hubs, thus a highly probable entrance point of this new lineage of *H. mabouia* into the island. Yet, the exact geographic origin of these animals is unclear as this lineage is genetically uniform and widespread across Central and South America. Likewise, since Brazil harbours both clades detected in Madeira (fig. 1), it precludes to disentangle if the Madeiran population of *H. mabouia* results from a single (i.e. same origin) or multiple introductions (i.e. distinct origins). At least in part of the Brazilian range of *H. mabouia* both lineages occur in sympatry (Rio Grande do Norte State; Carranza and Arnold, 2006). As such, the most parsimonious hypothesis would be that the specimens from Madeira have a Brazilian source, which would imply a single colonization event. However, this is a mere hypothesis that although plausible, cannot be confirmed by our results.

During our fieldwork we observed that *H. mabouia* is now well established in Madeira and that over the last 20 years the species has expanded from Funchal, where it was first detected (Jesus et al, 2002), to Câmara de Lobos. In both localities we observed numerous immature individuals and pregnant females, suggesting a healthy population. Yet, our observations highlight the need for an island-wide survey to assess the species’ distribution. Little is known also about the current range of another non-native gecko species occurring in Madeira, the Moorish gecko *Tarentola mauritanica*, which was first reported 8 km East of Funchal in 1993 (Báez and Biscoito, 1993). Yet, our 2020 fieldwork, shows that *T. mauritanica* has greatly expanded its distribution through the south cost of the island, and it can be found >20 km away from the locality where it was first recorded.

The negative consequences of introduced species, including non-native geckos, on native biodiversity are increasingly recognized (Doherty et al, 2016; Perella and Behm, 2020). *Hemidactylus mabouia* has been observed to compete with numerous species of diurnal and nocturnal geckos, including *H. angulatus* in Cameroon (Böhme, 1975), *Gymnodactylus darwinii* in Brazil (Teixeira, 2002; Zamprogno and Teixeira, 1997), *Phyllodactylus martini* and *Gonatodes antillensis* in Curaçao and Bonaire (Van Buurt, 2006), and *Gonatodes vittatus* and *Thecadactylus rapicauda* in Venezuela (Fuenmayor et al, 2005). Furthermore, *H. mabouia* also preys on a wide diversity of arthropods (Iturriaga and Marrero, 2013), potentially impacting native arthropod taxa in the species’ non-native range. So far, no study has addressed any potential effects of *H. mabouia* on the native fauna of Madeira. Yet, the island is home to a diverse arthropod fauna with numerous narrow range endemic species (Boieiro et al, 2014), which can be at risk as they might become part of the diet of the introduced geckos.

Over the last three decades the terrestrial reptile fauna of Madeira has increased from one, the endemic Madeira wall-lizard *Teira dugesii*, to at least four species due to the arrival and establishment of the geckos *T. mauritanica* (Báez and Biscoito, 1993) and *H. mabouia* (Jesus et al 2002) and of the brahminy blind snake *Indotyphlops braminus* (Jesus et al, 2013). The Mediterranean chameleon *Chamaeleo chamaeleon*, the Peters’s Rock Agama *Agama picticauda* (Wagner et al, 2012) and the Cape Verdean skink *Chioninia fogoensis* (Clemens and Allain, 2020) have also been reported to the island, however, the existence of established populations pends confirmation. Still, the rate of arrival and establishment is alarming and as such, local authorities should be vigilant regarding new arrivals and act to prevent their establishment as soon as they are detected. Likewise, efforts should be made to avoid that the already established species spread to the other islands of the archipelago and to the nearby Selvagens Archipelago, home to the endemic Selvagens gecko *Tarentola boettgeri bischoffi*.

Overall, our study uncovered unexpectedly high genetic diversity for *H. mabouia* in Madeira, but the pattern of phylogenetic relationships prevents us to ascertain if the island’s population results from a single or multiple introductions. Still, from a conservation point of view, *H. mabouia* and other introduced species in Madeira deserve careful monitoring, and phylogenetic studies such as the one presented here can offer important insights regarding the geographic origins of introduced taxa, a key aspect for the implementation of action plans to stop the flow of non-native species.

## Acknowledgments

CR is supported by a postdoctoral contract from FCT, Portugal (DL57/2016/CP1440/CT0005) and RR by a postdoctoral fellowship from ARDITI – Madeira’s Regional Agency for the Development of Research, Technology and Innovation (M1420-09-5369-FSE-000002). Field and lab work were supported by the project PTDC/BIA-EVL/27958/2017 (to CR) from FCT. The authors would like to acknowledge Angelica Crottini for providing samples of *H. mercatorius*. Specimens’ collection and tissue sample transport were performed under the permits 05/IFCN/2020 and 03/IFCN/2020 provided by the Forests and Nature Conservation Institute from the Madeiran Autonomous Region.

